# Inhibition of Striatal-Enriched Protein Tyrosine Phosphatase (STEP) Activity Reverses Behavioral Deficits in a Rodent Model of Autism

**DOI:** 10.1101/2020.04.17.047597

**Authors:** Manavi Chatterjee, Priya Singh, Jian Xu, Paul J. Lombroso, Pradeep K. Kurup

## Abstract

Autism spectrum disorders (ASDs) are highly prevalent childhood illnesses characterized by impairments in communication, social behavior, and repetitive behaviors. Studies have found aberrant synaptic plasticity and neuronal connectivity during the early stages of brain development and have suggested that these contribute to an increased risk for ASD. STEP is a protein tyrosine phosphatase that regulates synaptic plasticity and is implicated in several cognitive disorders. Here we test the hypothesis that STEP may contribute to some of the aberrant behaviors present in the VPA-induced mouse model of ASD. *In utero* VPA exposure of pregnant dams results in autistic-like behavior in the pups, which is associated with a significant increase in the STEP expression in the prefrontal cortex. The elevated STEP protein levels are correlated with increased dephosphorylation of STEP substrates GluN2B, Pyk2 and ERK, suggesting upregulated STEP activity. Moreover, pharmacological inhibition of STEP rescues the sociability, repetitive and abnormal anxiety phenotypes commonly associated with ASD. These data suggest that STEP may play a role in the VPA model of ASD and STEP inhibition may have a potential therapeutic benefit in this model.

## 1. Introduction

Autism spectrum disorder (ASD) is a group of highly prevalent heterogeneous disorders with significant social and economic burden [1]. It is a developmental disorder characterized by social communication deficits in addition to repetitive behaviors and interests [2]. Despite its high prevalence, the molecular mechanisms contributing to ASD is not understood [3]. Deciphering the complex pathophysiology of ASD will have a major impact on the development of potential therapeutics to improve quality of life in patients as well as alleviate the burden to families.

In an effort to address these issues, several genetic and environmentally induced rodent models were developed [4]. Among these, the valproic acid (VPA) rodent model is widely used. VPA is a teratogen and epigenetic modifier that is associated with a higher incidence of autism in humans, as suggested by epidemiological data [5, 6]. VPA models have relevance in human ASDs based on population studies and phenocopy several brain abnormalities and behavioral deficits present in human autism [7, 8]. The VPA rodent model of autism is widely used because of its relevance to human autism, its construct and face validity, and its ability to cause structural and behavioral deficits similar to those found in human autism [7].

STriatal-Enriched protein tyrosine Phosphatase (STEP) is a brain-enriched protein tyrosine phosphatase present in striatum and other brain regions [9, 10]. STEP normally opposes the development of synaptic strengthening through its ability to dephosphorylate and inactivate a number of synaptic proteins that include several protein kinases, ERK1/2, Pyk2, and Fyn [11–14]. In addition, STEP dephosphorylation of the ionotropic glutamate receptor subunits GluN2B and GluA2 results in internalization of NMDA and AMPA receptor complexes [15, 16]. In addition, STEP also plays a role in spine dynamics by dephosphorylating SPIN90 [17, 18].

The regulation of key synaptic proteins explains the role of STEP in several disorders. STEP levels are increased in Alzheimer’s disease [19], schizophrenia [20], and Parkinson’s disease [21], and in neurodevelopmental disorders such as fragile X syndrome [22]. Several pathways, including glutamate, dopamine and brain derived neurotrophic factor (BDNF) signaling regulate STEP activity [9, 23], and some of these signaling pathways are also implicated in ASD [24–26].

Previous studies have demonstrated that the expression of STEP is developmentally regulated in different brain regions from embryonic stage to adulthood [27]. However, whether STEP contributes to the pathophysiology of some ASD models has not been tested. Here we test the hypothesis that STEP plays a role in the VPA-induced mouse model of ASD. In addition, we evaluate whether a small molecule STEP inhibitor, TC-2153, reverses behavioral deficits commonly associated with autistic phenotypes.

## 2. Materials and methods

### 2.1 Animals and drug treatments

All protocols were approved by the Yale University Institutional Animal Care and Use Committee and strictly adhered to the NIH Guide for the Care and Use of Laboratory Animals. Time pregnant female C57/Bl6 mice were injected with either saline or VPA (Sigma-Aldrich, 600mg/kg, subcutaneous) on E12.5. The pregnant mice were housed singly and monitored until the pups were born. The pups were weaned at P22 and housed in groups of 2–5 in standard vented-rack cages in a 12:12 h light: dark cycle with food and water available *ad libitum*. Male pups were either sacrificed for biochemistry (cohort 1) or tested for behaviors (cohorts 2 and 3) (Figure 1).

**Figure 1:**
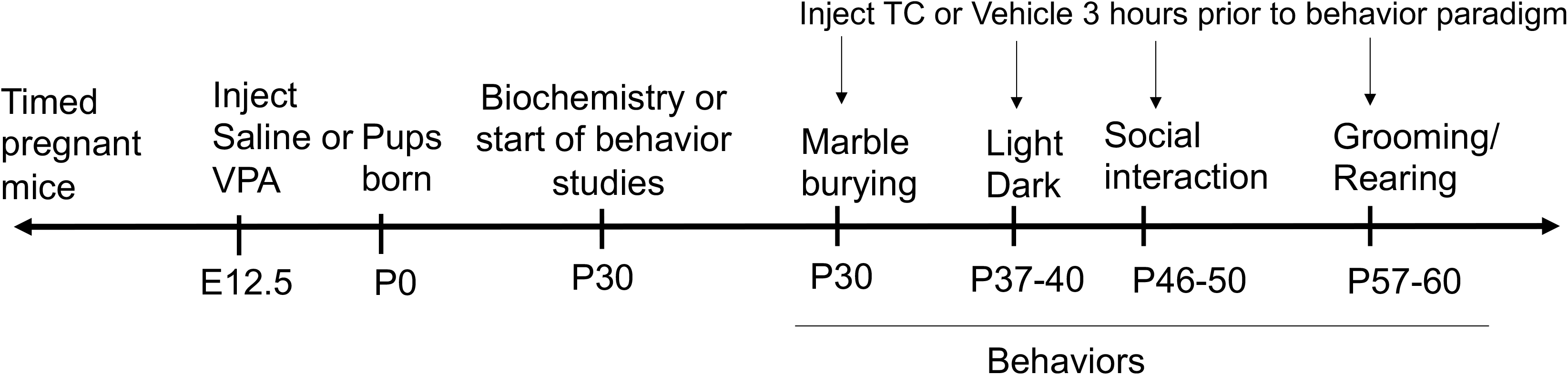
Timeline of drug injections and behavior experiments. Timed-pregnant mice were injected with VPA (600 mg/kg, s. c.) or saline at embryonic day 12.5 (E12.5). A group of mice were sacrificed at postnatal day 30 (P30) for biochemical analyses (Fig 3). Other mice were tested in behavioral studies (Figs 2 and 4) at the indicated time points. Some mice also received TC-2153 (i. p.) or vehicle administration 3 h prior to behavioral tests.

8-(tri-fluoromethyl)-1,2,3,4,5-benzopentathiepin-6-amine hydrochloride (TC-2153 (Sigma-Aldrich)) or vehicle (5% DMSO in saline) was administered through intraperitoneal injection at 10 mg/kg three hours prior to testing paradigms, as these parameters allows for adequate brain penetration [28].

### 2.2 Behavior tests

Animals were tested in the same order in behavioral tasks that included the marble burying, light/dark box, the three-chambered social choice task and grooming/rearing behaviors with a 7-10-day interval between tests (Figure 1). Drug treatments were assigned randomly on each test day. Behavior analysis was done using the ANY-maze tracking software to automate behavioral testing.

#### 2.2.1 Light/dark box

A plexiglass light/dark box (containing two equally-sized chambers joined by a small opening, where one side was clear and the other was black as well as covered with a black lid) was used to examine the aversion of mice to a brightly lit (i.e., anxiogenic) environments [29, 30]. Mice were placed in the light side of the light/dark box and were allowed to freely explore for 10 min. The time spent in each side were recorded using ANY-maze.

#### 2.2.2 Marble burying

Mice were allowed to acclimate individually in a cage filled with 4 cm of fresh bedding for 15 minutes. Mice were then removed from the cage and 12 black marbles were placed equidistant in a 3×4 arrangement. Mice were then returned to the same cage for 30 minutes. Following the 30-minute testing period, the number of marbles that were more than two-thirds covered with bedding were counted by a single blinded experimenter. After each testing period, the cage was cleaned with 70% ethanol, left to dry for 5 minutes, wiped down with a clean paper towel, and fresh bedding was then added. On completion, an overhead picture was captured. The percentage of each visible marble was quantified using ImageJ (version 1.43u; NIH). For each marble, the visible area was measured and compared with the area of a completely visible marble (control = 100% visible). If a marble was visible ≤60% of the control marble (i.e. buried at least 40%) were considered buried.

#### 2.2.3 Three-chambered social choice task

Sociability was evaluated by using the three-chambered social choice task, wherein mice were habituated in an opaque, rectangular plexiglass box divided into three equal compartments, each with a small entryway. Post habituation, mice were given the option to socialize with a conspecific stranger mouse enclosed under an inverted wire cup on one side, or a cup alone on the other side. We recorded the activity in close proximity (front 25% of the mouse within 1-inch margin around the cup) to the cups/mice in each chamber using the Any-maze software.

#### 2.2.4 Grooming

We analyzed repetitive behaviors in mice by scoring self-grooming time as previously described [31]. Each mouse was placed in a clean empty plastic cage for a 10-minute acclimation period without cage bedding (to avoid cage bed digging, a potentially competitive repetitive behavior). Following acclimation, mice were scored for cumulative time spent self-grooming and rearing episodes during a 10-minute testing period using a sideways digital camera. Between each testing period, the cage was cleaned with 70% ethanol, left to dry for 5 minutes, and wiped down with a clean paper towel.

### 2.3 Subcellular fractionation and Western blot analysis

Cortical tissue were dissected from saline or VPA exposed mice. Tissue was homogenized in buffer containing (in mM): 10 Tris base, pH 7.6, 320 sucrose, 150 NaCl, 5 EDTA, 5 EGTA, 1 NaF, 1 Na_3_VO4, 1 DTT, and protease inhibitor mixture (Roche). Homogenates were centrifuged at 800 × g to remove nuclei and large debris (P1), and crude synaptosomal membrane (P2) fractions were prepared by centrifugation of S1 at 9200 × g for 15 min. These fractions were protein estimated using BCA kit (Thermo Fischer). Proteins (30 μg) were loaded onto 8% (wt/vol) SDS/PAGE gels and transferred to nitrocellulose membranes, blotted with primary antibodies overnight at 4° C, followed by incubation with secondary antibody. Bands were visualized by chemiluminescence, using a G:BOX system with GeneSnap image program and quantified using Image J 1.33 (NIH).

### 2.4 Statistical analysis

All data are represented as mean ± SEM. Statistical differences were determined using student-t test using GraphPad Prism software. P<0.05 was considered significant.

## 3. Results

### 3.1 Administration of VPA *in utero* induced ASD phenotypes in pups

We first characterized the behavioral responses that are key to ASD phenotypes. We tested one-month old saline and VPA-exposed pups to light dark, marble burying, grooming and social interaction tasks. In agreement with previous papers [32, 33], VPA-exposed mice showed a significant increase in time spent in the light chamber of the box (t=2.172, df=17, p<0.05) in comparison to saline-exposed mice (Fig 2A). VPA-exposed mice also buried significantly more marbles (t=3.845, df=17, p<0.05, Fig 2B) and spend more time grooming (t=2.668, df=17, p<0.05, Fig 2C) compared to the saline-exposed controls. Rearing time was significantly reduced in these mice (t=2.462, df=17, p<0.05, Fig 2D). Furthermore, in the three-chambered task of social interaction, VPA-exposed mice spend significantly less time in the chamber containing a stranger mouse (t=2.245, df=36, p<0.05, Fig 2E). They display no preference for the social chamber whereas the control mouse shows a significant preference (t=1.363, df=16, Fig 2F) towards the chamber containing the mouse versus the empty chamber. VPA-exposed mice also spend significantly less time in close proximity to the stranger mouse (t=2.24, df=36, p<0.05, Fig 2G).

**Figure 2:**
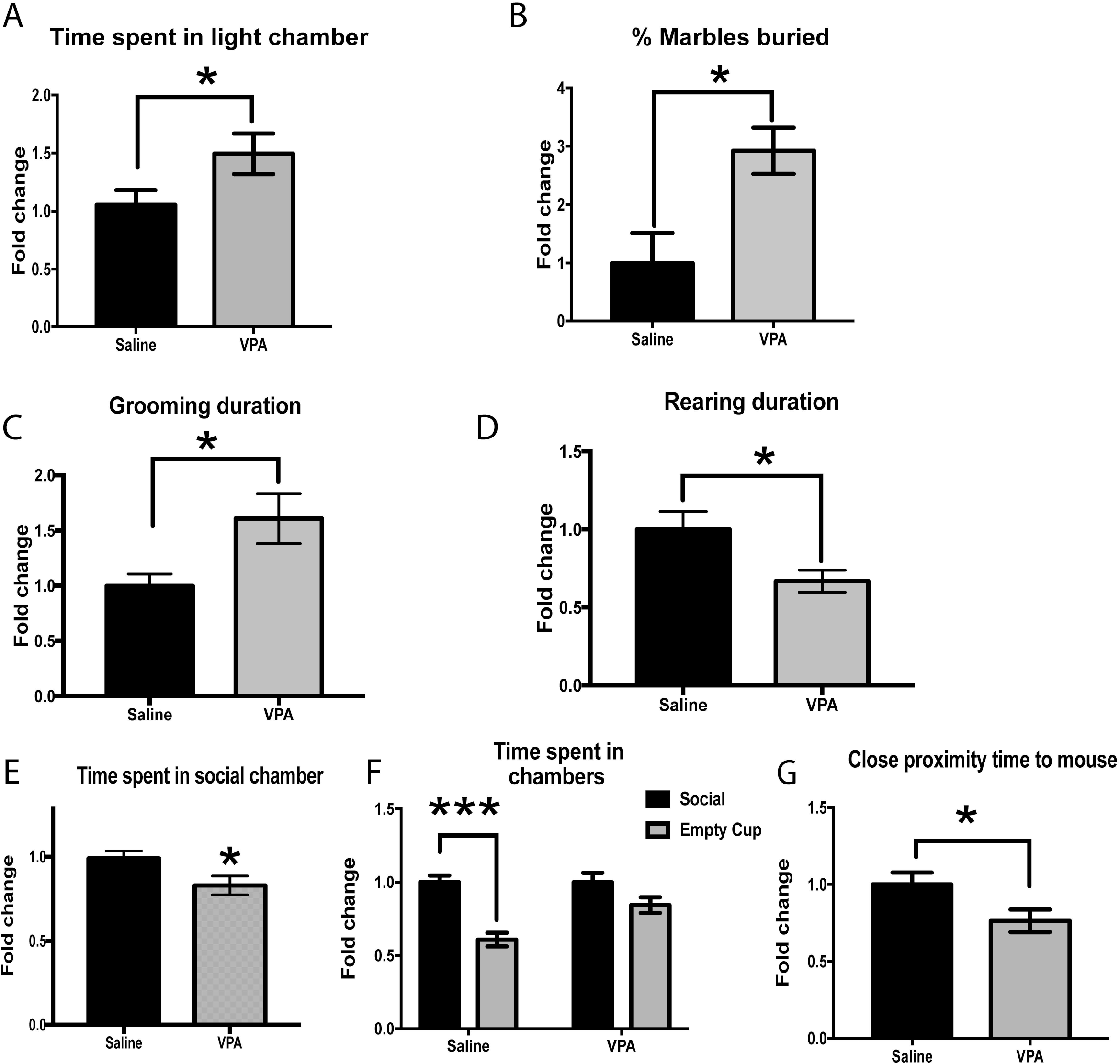
*In utero* VPA exposure in mice leads to ASD phenotype in mice. In utero VPA or saline exposed mice were subjected to behavioral tasks that are common to ASD in mice. VPA mice spent more time in the light chamber in the light dark box (A) and buried more marble in the marble burying task (B). VPA exposed mice also showed increased grooming (C) and reduced rearing (D) duration. In the social chamber task, they spent significant less time in the social chamber (E), show no preference for social versus empty chambers (F) as well as spend lesser time in close proximity to stranger mice (G) (\**p* < 0.05, Student t-test, n=8-12 per group).

### 3.2 Administration of VPA *in utero* leads to increased STEP61 activity in pups

Previous studies show disrupted synaptic development in pre-frontal cortex after VPA treatment in rodents [34, 35]. We therefore determined total STEP61 levels in pre-frontal cortex after pre-natal VPA-exposure in mice at one-month age by immunoblot analysis. There was a significant (t=4.204, df=5, p<0.01) increase in STEP61 levels in mice exposed to VPA prenatally and a decrease in the tyrosine phosphorylation of the STEP substrates, GluN2B (t=7.299, df=5, p<0.001), Pyk2 (t=8.032, df=4, P<0.01) and ERK(t=3.348, df=4, P<0.05) (Fig 3).

**Figure 3:**
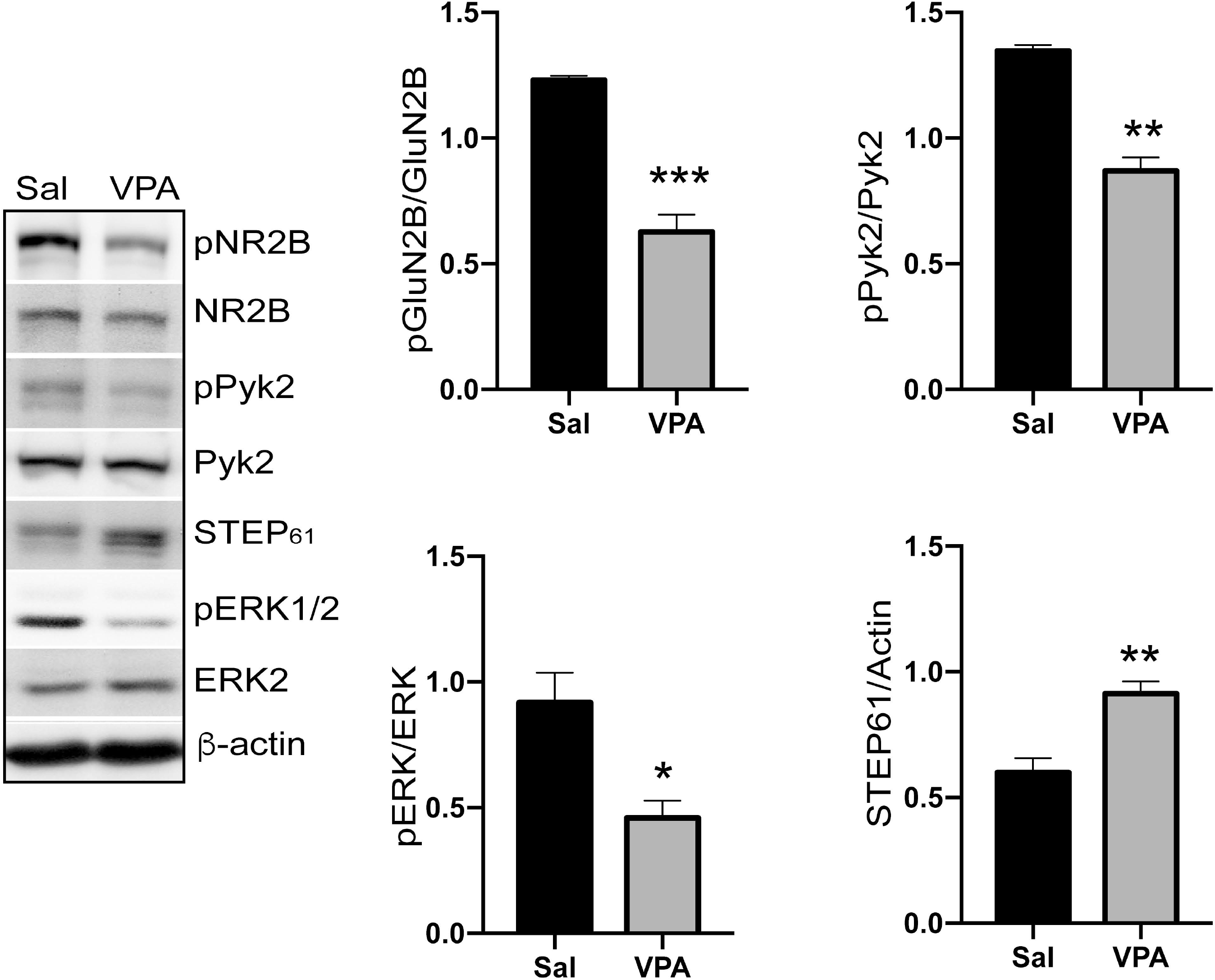
VPA increases STEP61 expression in cortical synaptosomal membrane fractions in mice. Immunoblot and quantification of STEP61 protein levels and phosphorylation levels of STEP substrates (GluN2B, Pyk2 and ERK1/2) in one-month old mice after prenatal exposure to saline or VPA in E12.5 mice. (\**p* < 0.05, \*\**p* < 0.01, \*\*\**p* < 0.001, Student t-test, n=3-4 per group).

### 3.3 Inhibition of STEP rescues ASD phenotypes in VPA exposed mice

We next assessed the effects of the STEP inhibitor TC-2153 on the behavioral phenotypes induced by VPA exposure. TC-2153 administration rescued the abnormal anxiety phenotype by significantly reducing the time spent in light chamber as compared to the vehicle treated VPA exposed mice (t=2.288, df=14, p<0.05, Fig 4A). Administration of TC-2153 also reduced the repetitive behaviors by reducing the number of marbles buried as compared to the vehicle treated group (t=2.481, df=14, p<0.05, Fig 4B). TC-2153 treatment also reduced the duration of grooming (t=3.267, df=14, p<0.01 Fig 4C); however, it had no significant effects in rearing duration (t=0.6669, df=14, Fig 4D). In the sociability tasks, TC-2153-treated mice (exposed to VPA) spent significantly more time in the social chamber (t=3.391, df=27, p<0.01, Fig 4E). They also showed a significant preference (t=7.405, df=11, p<0.001) towards the social chamber versus the empty cup compared to the vehicle treated VPA mice (t=7.405, df=11, p<0.001, Fig 4F). Furthermore, the time spent in close proximity was also increased in the TC-2153 mice as compared to vehicle-treated VPA controls (t=3.034, df=27, p<0.01, Fig 4G).

**Figure 4:**
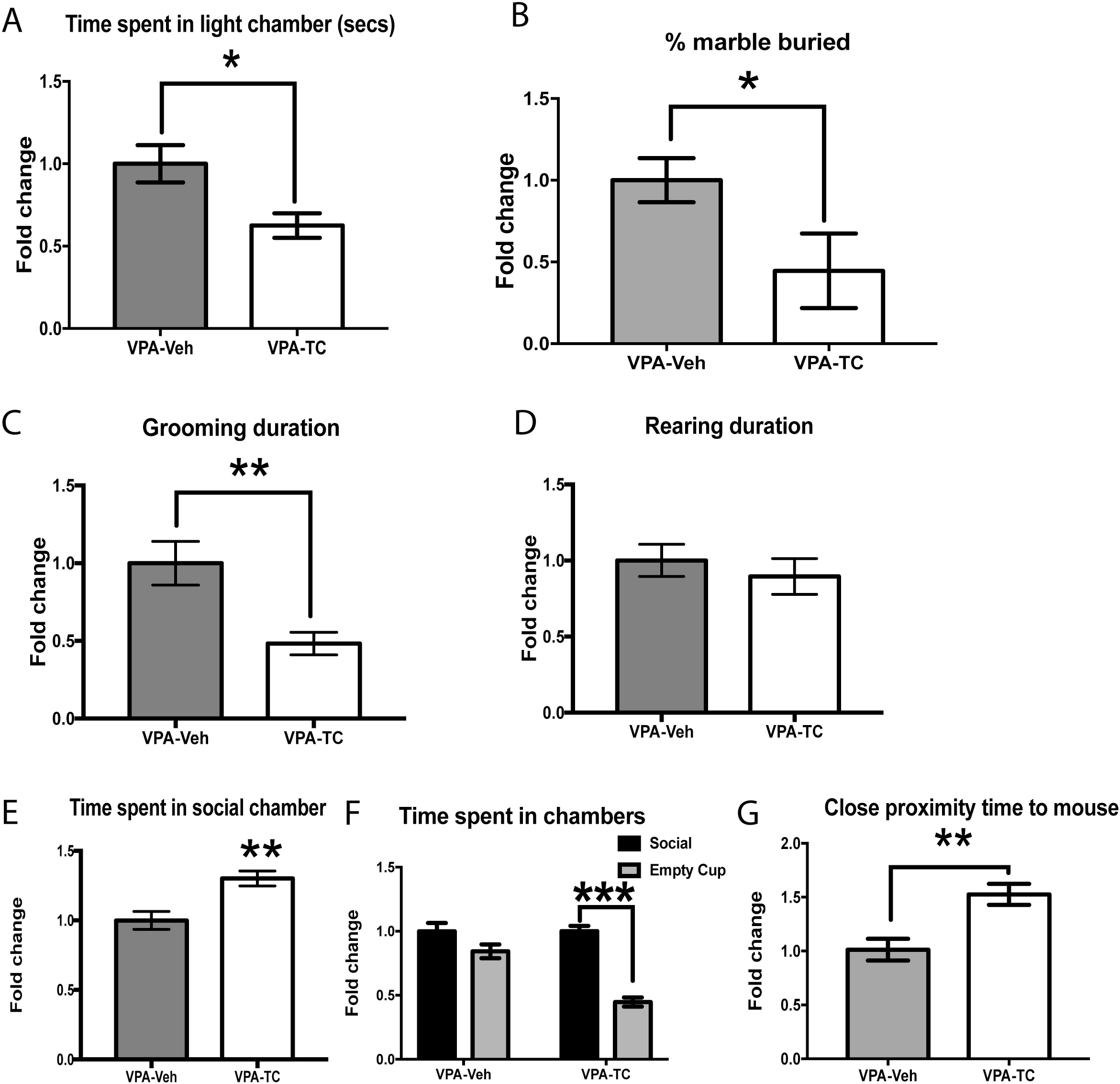
STEP inhibitor TC-2153 attenuates ASD in VPA exposed mice. VPA exposed mice were pretreated with TC-2153 or vehicle, three hours prior to behavioral testing. TC-2153 treatment significantly reduced the time spent in light chambers in light dark task (A). TC-2153 also showed reduction in marble burying behaviors (B) as well as grooming duration (C) in the VPA mice as compared to vehicle treated mice. No significant differences were observed in the rearing duration (D). In the sociability task, TC-2153 treated VPA mice spent significantly more time in the social chamber (E) and showed increased preference for the social chamber as compared to the empty chamber (F). TC-2153 treated VPA mice also spent more time in the close proximity to the stranger mouse as compared to the vehicle treated VPA mice (G) (\**p* < 0.05, \*\**p* < 0.01, \*\*\**p* < 0.001, Student t-test, n=8-12 per group).

## 4. Discussion

Autism Spectrum Disorder (ASD) is a neurodevelopmental disorder with known genetic, environmental, and epigenetic etiologies [2]. A major hypothesis in the field is that ASD is characterized by aberrant neuronal trajectory during early stages of development, which leads to subsequent functional alterations in neuronal regional connectivity [36, 37].

We explored the role of STEP in the VPA-exposed mice model of autism. *In utero* exposure of VPA leads to behavioral abnormalities in rodents that are similar to some ASD phenotypes [4, 8]. In agreement with earlier studies, we found that VPA-exposed mice showed reduced social preference, increased repetitive behaviors, and abnormal anxiety behaviors (Fig 2) [5, 32, 38–40]. STEP61 levels were upregulated in the prefrontal cortex regions in mice exposed prenatally to VPA (Fig 3). The increase in STEP correlates with a decrease in the tyrosine phosphorylation of its substrates (ERK, Pyk2, GluN2B) (Fig 3). Pharmacological reduction of STEP activity rescued the behavioral phenotypes induced by *in utero* VPA-exposure (Fig 4).

The rationale for studying STEP stems from its role in fragile X syndrome (FXS), which shares a significant overlap with ASD [41]. Increased STEP activity has been observed in *Fmr1* KO mice. Fragile X mental retardation protein (FMRP) directly interacts with STEP mRNA [42] and regulates STEP expression levels [22]. *Fmr1* KO mice have higher levels of STEP and both genetic reduction and pharmacologic inhibition of STEP reversed some of the phenotypic deficits in the mouse model of FXS, including social impairments that were present [22, 43].

A similar increase in STEP61 expression is observed in the VPA-exposed pups. VPA being an epigenetic modulator and an inhibitor of histone deacetylase (HDAC), has been known to modulate the expression of multiple genes [44]. It could be possible that VPA alters the expression of STEP at a transcriptional level. Further studies are needed to test the hypothesis. Another possibility could be that the increase in STEP61 expression is due to reduction of BDNF expression [45], as reported in patients as well as animals exposed to VPA [5, 46]. Previous studies from the lab show that STEP and BDNF share a reciprocal relationship. BDNF/Trk B signaling pathways promotes the ubiquitination and degradation of STEP [23, 45] and thereby reduces STEP levels. In addition, STEP regulates BDNF expression through ERK/CREB pathway [47]. Moreover, the STEP inhibitor, TC-2153, increases BDNF gene expression [48].

STEP accumulation and overactivation may have several downstream effects. For instance, STEP expression is critical during the development of midbrain dopaminergic neurons and mediates D2-induced ERK activation [27]. Overactivation of STEP in VPA-exposed mice might lead to reduced ERK signaling in these neurons. Disruption in ERK signaling during this critical neurodevelopment period is known to cause social impairment, increased anxiety and grooming behaviors in mice later in adulthood [49]. Recent studies have also found higher dopamine turnover in VPA-exposed mice [50]. Since co-activation of D1 and D2 receptors are implicated in social deficits [51] and D2 receptors control DA release and neurotransmission [52, 53], it is possible that the reduction of D2/ERK signaling due to overactivation of STEP results in increased DA firing through co-activation of D1 and D2-R coactivation as a feedback mechanism. Moreover, STEP leads to the dephosphorylation of the GluN2B subunit of NMDA receptor, leading to the internalization of NMDA receptor complexes. Loss of glutamatergic proteins has been associated with social impairments in VPA-exposed mice [54].

In conclusion, we report that inhibition of STEP activity attenuates social and behavioral deficits in the VPA model of autism in mice. These pre-clinical experiments need to be expanded to further our understanding of the potential role of STEP in ASD.

## Acknowledgments

We thank Marija Kamceva for technical support and other laboratory members for helpful discussions.

## Author contributions

MC, PKK and PJL conceived of and designed the experiments. JX and PKK performed drug injections and biochemistry. MC and PS performed in vivo drug administration and behavior experiments. MC, PKK and PJL interpreted the data. MC, JX, PKK and PJL helped write the manuscript. All authors approved the final version of the submitted manuscript.

